# Serial sarcomere number is substantially decreased within the paretic biceps brachii in individuals with chronic hemiparetic stroke

**DOI:** 10.1101/2020.03.12.989525

**Authors:** Amy N. Adkins, Julius P.A. Dewald, Lindsay Garmirian, Christa M. Nelson, Wendy M. Murray

**Affiliations:** Department of Biomedical Engineering, Northwestern University, Evanston, IL, 60208; Department of Physical Therapy and Human Movement Sciences, Northwestern University Feinberg School of Medicine, Chicago, IL, 60610; Department of Physical Medicine & Rehabilitation, Northwestern University Feinberg School of Medicine, Chicago, IL, 60610; Shirley Ryan AbilityLab (formerly the RIC), Chicago, IL, 60610; Edward Hines, Jr. VA Hospital, Hines, IL, 60141; Department of Health, Human Function & Rehabilitation Sciences, George Washington University, Washington, DC, 20052; Department of Physical Therapy and Rehabilitation Science, University of Maryland School of Medicine, Baltimore, MD, 21201

**Keywords:** Muscle, Stroke, Sarcomere, Fascicle, Imaging

## Abstract

A muscle’s structure, or architecture, is indicative of its function and is plastic; changes in input to or use of the muscle alter its architecture. Stroke-induced neural deficits substantially alter both input to and usage of individual muscles. Here, we combined novel *in vivo* imaging methods (second harmonic generation microendoscopy, extended field-of-view ultrasound, and fat-supression MRI) to quantify functionally meaningful muscle architecture parameters in the biceps brachii of both limbs of individuals with chronic hemiparetic stroke and in age-matched, unimpaired controls. Specifically, serial sarcomere number and physiological cross-sectional area were calculated from data collected at three anatomical scales: sarcomere length, fascicle length, and muscle volume. Our data indicate that the paretic biceps brachii had ~8,500 fewer serial sarcomeres compared to the contralateral limb (p=0.0044). In the single joint posture tested, the decreased serial sarcomere number was manifested by significantly shorter fascicles (10.7cm vs 13.6cm; p<0.0001) without significant differences in sarcomere lengths (3.58μm vs. 3.54μm; p=0.6787) in the paretic compared to the contralateral biceps. No interlimb differences were observed in unimpaired controls, suggesting we observed muscle adaptations associated with stroke rather than natural interlimb variability. This study provides the first direct evidence of the loss of serial sarcomeres in human muscle, observed in a population with neural impairments that lead to disuse and chronically place the affected muscle at a shortened position. This adaptation is consistent with functional consequences (increased passive resistance to elbow extension) that would amplify already problematic, neurally driven motor impairments.

**SIGNIFICANCE STATEMENT:** Serial sarcomere number determines a muscle’s length during maximum force production and its available length range for active force generation. Skeletal muscle length adapts to functional demands; for example, animal studies demonstrate that chronically shortened muscles decrease length by losing serial sarcomeres. This phenomenon has never been demonstrated in humans. Integrating multi-scale imaging techniques, including two photon microendoscopy, an innovative advance from traditional, invasive measurement methods at the sarcomere scale, we establish that chronic impairments that place a muscle in a shortened position are associated with the loss of serial sarcomeres in humans. Understanding how muscle adapts following impairment is critical to the design of more effective clinical interventions to mitigate such adaptations and to improve function following motor impairments.

## INTRODUCTION

Three-fourths of the nearly 7 million stroke survivors currently living in the United States report substantial motor impairments that limit their independence in tasks of everyday life(1, 2). Damage to the corticofugal motor pathways and the resulting reliance on indirect corticoreticulospinal pathways following stroke alter neuronal input to contralesional muscles(3–6). Such changes include decreased voluntary neural drive (weakness or paresis)(7, 8) and increased involuntary neural drive (hypertonicity), causing the inability to fully activate and fully relax the muscle, respectively. Further, abnormal muscle coactivation patterns(9) also arise from the altered neuronal drive, commonly leading to abnormal limb synergies(10) or a loss of independent joint control(9, 11). For example, in the upper limb, the flexion synergy presents when the individual attempts to abduct their shoulder but involuntarily and concomitantly flexes all of their distal joints (elbow, wrist, and fingers)(9, 12). The flexion synergy limits the ability to combine shoulder abduction and extension of the limb, as needed to use the arm and hand to reach and acquire an object at a distance from the body. Thus, although stroke is primarily an injury of the brain, stroke-induced neural deficits substantially alter input to and use of the contralesional upper limb.

Skeletal muscle is a plastic tissue; changes to both the stimulus it receives and how it is used can alter its functional capacity (e.g.(13–19)). A muscle’s function, its ability to contract and produce force, can be delineated via its architecture(20–23). Specifically, optimal fascicle length (OFL) is the fascicle length at which the muscle generates its maximum force. OFL is a measure of the number of sarcomeres in series in the muscle, so it also provides a measure of the absolute range of lengths over which the muscle can actively generate force(21, 24). Physiological cross-sectional area (PCSA) is a correlate of the maximum isometric force a muscle can produce given maximum activation(25). These two critical muscle architectural features are calculated from more primary measures of a muscle’s structural anatomy that occur across different scales, including muscle volume, fascicle length, pennation angle (in pennate muscle), and sarcomere length (see Methods, Eqs. 4 and 5)(24, 26). Quantifying the adaptation of these two parameters to changes in use and stimulus can provide insight into the functional implications of such changes. For example, a series of classic studies in animal models demonstrate that when limbs were immobilized for an extended time, muscles immobilized at a joint angle that shortened muscle-tendon lengths (i.e., with origin-to-insertion distances that were decreased compared to resting length) lost serial sarcomeres (i.e., adapted such that OFL was shorter)(18, 27–29). Conversely, when immobilized at a joint posture that lengthened muscles, sarcomeres were added in series. These fundamental studies suggest that when *in vivo* length is chronically altered, a muscle’s architecture changes in a way that maximizes its function at the new length. Specifically, in the immobilization studies, the muscle’s length “ reoptimized” so OFL (the length at which the muscle generates its maximum force) occurred at the joint angle of immobilization.

Despite these important studies, it is unclear the extent to which this mechanism is expressed, *in vivo,* in human muscle. For example, at this point, only a single published study has calculated serial sarcomere number (i.e., measured both sarcomere length and fascicle length *in vivo*) under conditions in which a human muscle has chronically been placed at a *Λ*shortened position^32^. In this case, the calf muscles in children with cerebral palsy with equinus contractures (functionally shortened muscle-tendon units that are more resistant to stretch(30, 31)) severe enough to require surgical intervention did not “ re-optimize” to the shorter muscle lengths imposed by the participants’ chronically plantarflexed joint posture. Rather, the soleus muscles in these children had sarcomeres substantially longer (4.07μm(32)) than optimal length (2.70μm(33)). Unfortunately, identical measurements could not be collected in a control population because sarcomere length could not be measured in children who were not undergoing surgery. Sarcomere length measurements in living human subjects have traditionally been limited to biopsy or intraoperative studies, (i.e. during distraction surgeries for limb discrepancy(13), tendon transfer surgeries(34, 35), or muscle lengthening in children with cerebral palsy(30)) and only recently have become measurable via minimally or non-invasive techniques(36, 37). As a result, the authors estimated serial sarcomere number for control participants by combining fascicle length measures obtained from typically developing children via ultrasound with sarcomere lengths measured from adult cadavers via dissection. Based on these methods, the authors concluded the soleus in the children with CP had fewer sarcomeres in series than typically developing muscle(32). This pediatric study suggested that, instead of reoptimizing so that OFL occurred at the chronically plantarflexed ankle angle, the shortened muscle shifted to much longer sarcomere lengths, at which active force-generating capabilities are weaker and passive forces are greater.

Stroke-induced neural deficits lead to decreased use and a chronically more flexed resting posture of the paretic upper limb. Given the muscle adaptation observed following immobilization in animal models, as well as a general assumption that these type of muscle adaptations do occur in humans with severe neural impairments and may result in muscles that are both structurally weaker and stiffer, there is concern about functional consequences of muscle adaptation following stroke. Thus, several studies have sought to directly quantify *in vivo* muscle structure in this cohort. In general, *in vivo* medical imaging has provided strong evidence that many muscle anatomical structural parameters are different in the paretic limb compared to the contralateral limb. Specifically, shorter fascicles(38), as well as smaller pennation angles(39), muscle masses(40), volumes(41, 42), and anatomical cross-sectional areas (the area of muscle perpendicular to the line of action of the external tendons(43)) have been demonstrated in numerous thigh and shank muscles(41, 44). Similarly, in the upper limb, shorter fascicles(45, 46) and smaller muscle volumes(47) have been reported in various paretic muscles as compared to the contralateral side.

The primary weakness of existing studies that demonstrate differences in anatomical parameters in paretic muscles in chronic stroke is that none also simultaneously measured sarcomere length. Neither OFL nor PCSA is calculable in any existing study that describes muscle structure following chronic stroke because they do not report the corresponding measures of both a muscle’s sarcomere and fascicle lengths. Thus, insight into how the observed structural differences relate to muscle function in the paretic limb is limited. For example, shorter fascicles in the paretic limb relative to the contralateral limb have been widely demonstrated(38, 45, 46). One explanation of this observation in the elbow flexors could be that they experience conditions similar to muscle immobilization at shortened muscle-tendon lengths; the paretic upper limb is generally used much less than the non-paretic limb and it rests in a more flexed elbow posture(48). Thus, similar to immobilization studies in animal models (i.e.(18, 27, 29)), shorter fascicles in the paretic elbow flexors could result from a loss of sarcomeres in series. Functionally, this adaptation would indicate a decrease in the absolute range of lengths over which the muscle can generate active force(21). In addition, the animal studies demonstrated that muscles that lost sarcomeres in series exhibited a shift in the onset of passive force development to shorter lengths; the muscles that lost sarcomeres in series were also described as less “ extensible” (29). However, without a concomitant measure of the length of the muscle’s sarcomeres, the possibility that fascicle lengths measured in paretic muscle are shorter because there are fewer sarcomeres in series cannot be distinguished from the possibility that the paretic muscle has the same number of sarcomeres in series as in the contralateral limb, but its sarcomeres have adapted by shifting to shorter lengths. In this case, the paretic muscle would be capable of generating active force over the same range of lengths, but the muscle would operate over shorter sarcomere lengths throughout the joint’s range of motion. Based on basic muscle physiology, we would expect a muscle operating at shorter sarcomere lengths to generate smaller passive forces throughout the joint’s range of motion(43). A single study has reported *in vivo* measures of biceps brachii sarcomere lengths following chronic hemiparetic stroke (fascicle lengths were not measured); 2 of 4 stroke participants had longer sarcomeres in the paretic biceps compared to the non-paretic side, and 2 had shorter sarcomeres(49). Thus, the implications for OFL, the range of lengths over which the muscle can generate active force, and the passive forces the muscle produces over the joint’s range of motion, remain unclear. These distinct possibilities for sarcomere adaptation would also have different effects on maximum isometric force capacity. While a decrease in muscle volume is regularly reported in paretic limbs(41, 47), PCSA is the architectural correlate of force-generating capacity, and it is calculated using the ratio between muscle volume and OFL. Critically, stroke alters both neural input(5–7) to and use(8, 9, 11, 12) of the paretic limb. Thus, it is also possible that paretic muscle adapts in an unexpected manner post-stroke; animal models of immobilization limited the use of the limb and altered muscle length but did not involve neural injury.

In this study, we quantify multiscale muscle parameter differences, *in vivo,* in the biceps brachii of the paretic and contralateral limbs of individuals with chronic hemiparetic stroke and in both limbs of a group of age-matched individuals with no history of musculoskeletal or neurological diseases or injuries to the upper limb. Specifically, sarcomere length, fascicle length, and muscle volume are measured from images obtained using second harmonic generation microendoscopy, extended field-of-view ultrasound, and fat suppression magnetic resonance imaging, respectively. Our most prominent finding was that the paretic biceps of individuals with chronic hemiparetic stroke has fewer sarcomeres in series (i.e., shorter OFL) compared to the contralateral muscle. In the limb posture we evaluated, this result was manifested by systematically shorter fascicles in the paretic muscle without significantly different sarcomere lengths from the non-paretic, contralateral biceps. Our data provide the first direct evidence of muscle adaptation involving the loss of serial sarcomeres in living human subjects and is observed in a population with neural impairments that chronically place the affected muscle at a shortened position. This muscle architectural difference post chronic hemiparetic stroke is consistent with functional consequences that would amplify the already problematic neural driven motor impairments (i.e. the ability to reach away from the body to grab an object).

## RESULTS

### SSN, Fascicle length, and Sarcomere Length

Relative to the contralateral, non-paretic limb, the paretic biceps brachii of individuals with chronic hemiparetic stroke had fewer sarcomeres in series; this inter-limb difference was not observed in participants without stroke. Specifically, there was a statistically significant difference in overall SSN between paretic and non-paretic limbs (**p = 0.0044**) with an estimated mean SSN in the paretic muscle of 30,152 ± 2,163 (95% CI, 25,038 to 35,267) compared to 38,697 ± 2,162 (95% CI, 33,584 to 43,810) serial sarcomeres in the non-paretic muscle (Fig 1). We also observed significantly shorter biceps fascicles (**p < 0.0001**) in these participants’ paretic limbs; mean fascicle lengths on the paretic side were shorter (10.66 ± 0.61 cm; 95% CI, 9.46 to 11.85 cm) than non-paretic fascicle lengths (13.59 ± 0.61 cm; 95% CI, 12.40 to 14.79 cm) in the single joint posture we tested (Fig. 2A). There was no significant difference in sarcomere length in this posture (p = 0.6787; paretic 3.58 ± 0.08 μm 95% CI, 3.42 to 3.73 μm; non-paretic 3.54 ± 0.08 μm 95% CI, 3.39 to 3.70 μm) (Fig. 2B). For the individuals who did not have a stroke, there were no observed interlimb differences in SSN (p = 0.2463; dominant 40,102 ± 1,451 95% CI, 37,253 to 42,951; non-dominant 39,545 ± 1,453 95% CI, 36,691 to 42,398), fascicle length (p = 0.0790; dominant 14.32 ± 0.27 cm 95% CI, 13.78 to 14.86 cm; non-dominant 14.11 ± 0.27 cm 95% CI, 13.57 to 14.65 cm), or sarcomere length (p = 0.9423; dominant 3.58 ± 0.07 μm 95% CI, 3.44 to 3.72 μm; non-dominant 3.59 ± 0.07 μm 95% CI, 3.45 to 3.73 μm) (Fig 2–3). The relationship between Fugl-Meyer Assessment (FMA) and percent difference in fascicle length yielded a modest linear relationship (R^2^ = 0.5953) which was significant (**p = 0.0249**) (Fig. 3A); the relationship between FMA and percent difference in SSN was weaker (R^2^ = 0.4676; p = 0.0615) (Fig. 3B).

**Figure 1:**
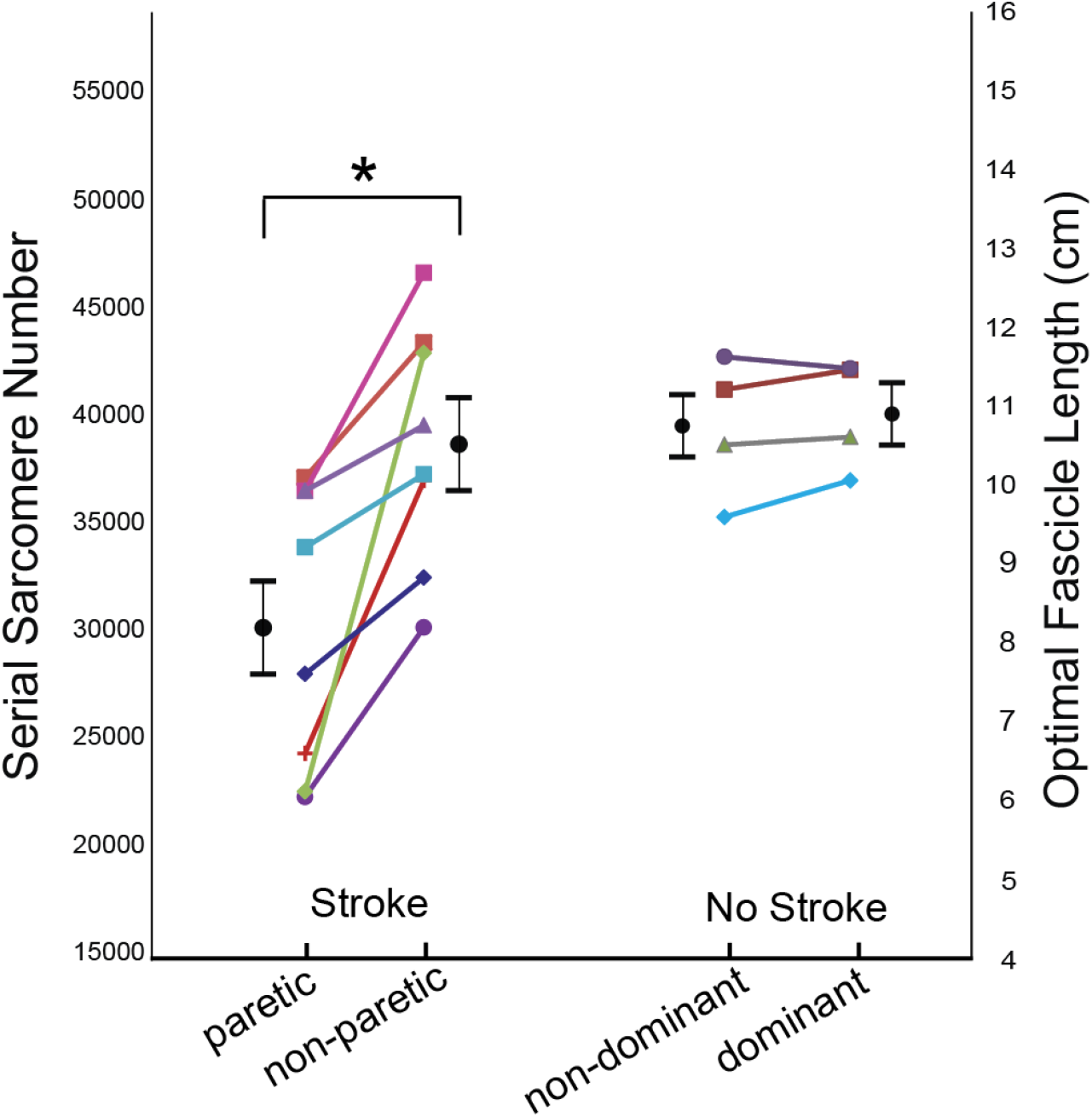
Serial Sarcomere Number/ Optimal Fascicle Length. Data showing interlimb differences in serial sarcomere number and, proportionally, optimal fascicle length for all participants who had a stroke and the age-range matched controls (no stroke). Each participant is represented by a different shape/color and each individual’s limbs are connected by a line. Black circular points and error bars which are offset from individual participant data, represent mean and standard deviations estimated from the generalized mixed effects model. The star (*) indicates a significant interlimb difference (p<0.05).

**Figure 2:**
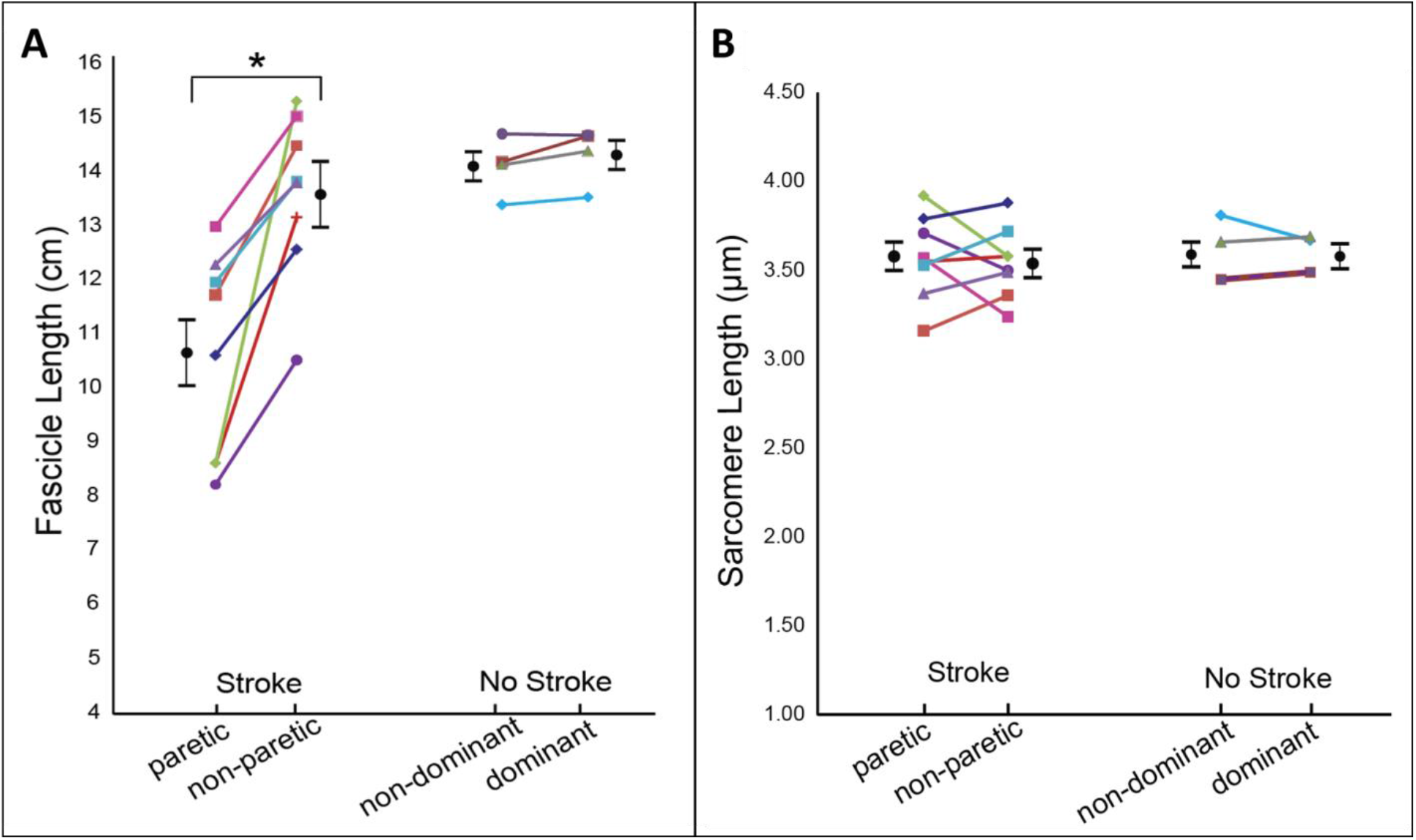
Fascicle and Sarcomere Length. Graphs displaying mean sarcomere length (A) and fascicle length (B) measurements from both limbs and in all participants in the stroke and control (no-stroke) group. Each participant is represented by a different shape and or color. Each individual’s limbs are connected by a solid line with the exception of one healthy individual whom had the same average sarcomere length on both limbs as another healthy individual (dashed line to enable visualization of both healthy individuals). Black circular points and error bars which are offset from individual participant data, represent mean and standard deviations estimated from the generalized mixed effects model. The star (*) indicates a significant interlimb difference (p<0.05).

**Figure 3:**
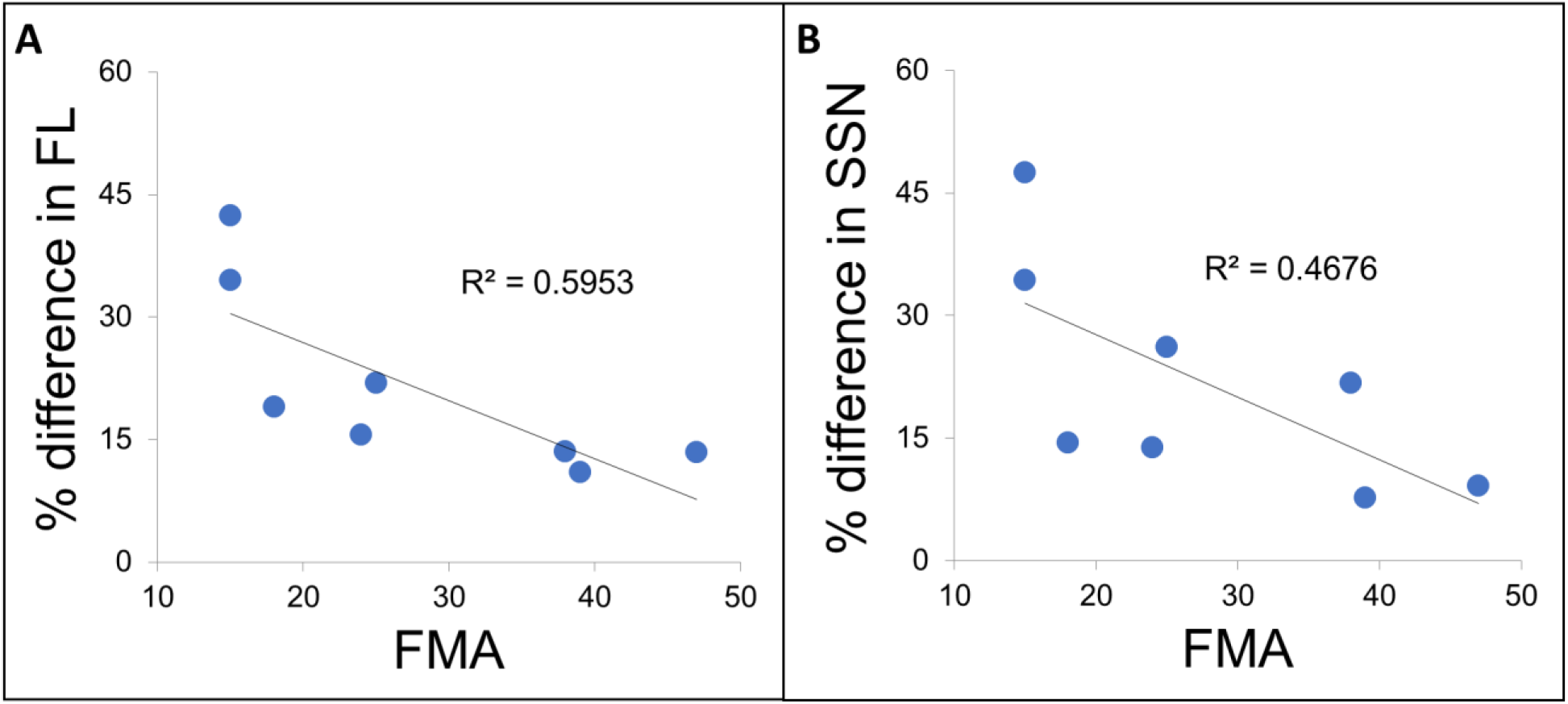
Relationship between change in muscle parameters and clinical assessment. Graphs showing the relationship between percent difference in serial sarcomere number (A) and percent difference in fascicle length (B) versus the Fugl-Meyer Assessment clinical impairment score. There is a trend toward a greater difference in OFL as the impairment level increases (Fugl-Meyer becomes smaller number).

**Table 1:** Demographic information of individuals who participated in this study.

|  | Gender | Age* | Paretic | Years Post Stroke* | FMA |
| --- | --- | --- | --- | --- | --- |
| 1 | F | 65 | R | 32 | 25 |
| 2 | F | 65 | R | 11 | 15 |
| 3 | F | 63 | L | 15 | 24 |
| 4 | M | 48 | L | 8 | 18 |
| 5 | M | 60 | R | 7 | 15 |
| 6 | M | 72 | R | 22 | 38 |
| 7 | M | 62 | L | 5 | 47 |
| 8 | M | 44 | R | 5 | 39 |
|  | Gender | Age* | Dominant |  |  |
| 9 | F | 53 | R |  |  |
| 10 | F | 64 | R |  |  |
| 11 | M | 62 | R |  |  |
| 12 | M | 67 | R |  |  |

\*Age for all participants and years post-stroke for participants with stroke (Subjects 1-8) are reported as of the time at which the experimental data collection for that participant was completed.

### PCSA and Muscle Volume

There was a substantial difference in muscle volume and a modest difference in PCSA in the paretic biceps brachii in participants with chronic hemiparetic stroke as compared to the contralateral limb. Muscle volume was significantly smaller on the paretic side (**p = 0.0127**; paretic 105 ± 21 cm^3^ 95% CI, 55 to 154 cm^3^; non-paretic 148 ± 21 cm^3^ 95% CI, 99 to 197 cm^3^); however, one stroke participant had a larger muscle volume on the paretic side compared to the non-paretic side (Fig. 4A). This individual also had the smallest volume among all participants’ non-paretic or dominant limbs. On average, post-stroke participants had larger PCSAs in their non-paretic limb (p = 0.2075; paretic 12.59 ± 1.71 cm^2^ 95% CI, 8.55 to 16.64 cm^2^; non-paretic 13.72 ± 1.71 cm^2^ 95% CI, 9.66 to 17.76 cm^2^) (Fig 4B). We did not observe interlimb differences in either volume (p = 0.7743; dominant 125 ± 40 cm^3^ 95% CI, −3 to 253 cm^3^; non-dominant 124 ± 40 cm^3^ 95% CI, −3 to 253 cm^3^) or PCSA across limbs (p = 0.5561; dominant 11.38 ± 3.45 cm^2^ 95% CI, 4.60 to 18.16 cm^2^; non-dominant 11.52 ± 3.45 cm^2^ 95% CI, 4.74 to 18.29 cm^2^) in the participants who had not had a stroke (Fig 4). In the seven stroke participants with smaller biceps volumes in the paretic limb, normalized interlimb differences in PCSA were smaller than those in volume due to fewer sarcomeres in series (Fig 5).

**Figure 4:**
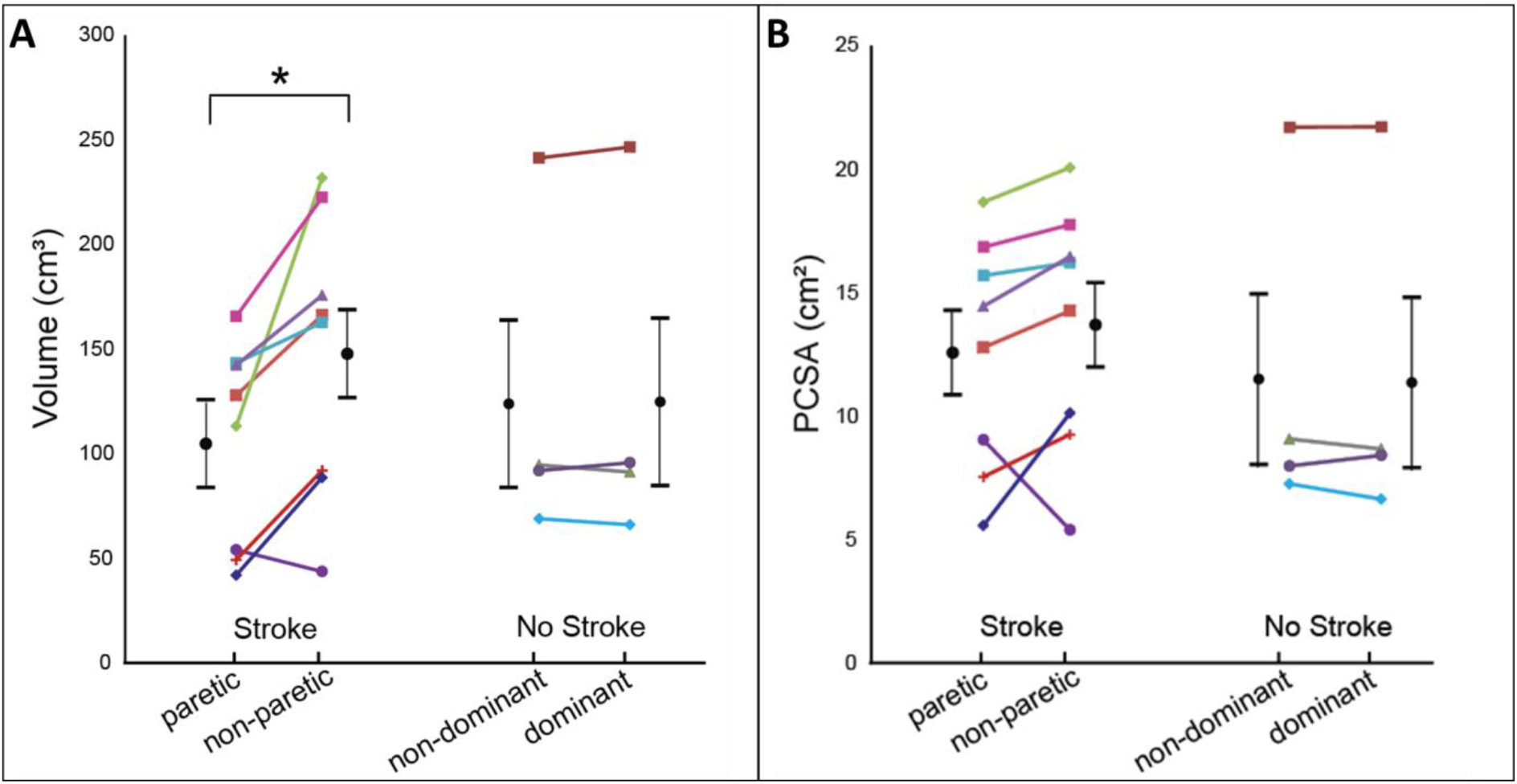
Muscle Volume and PCSA. Data for muscle volume with intramuscular fat removed (A) and calculated PCSA (B) for both biceps braciii of all participants who had a stroke and the age matched controls (no stroke). Each participant is represented by a different shape and or color and each individual’s limbs are connected by a solid line. Black circular points and error bars which are offset from individual participant data, represent mean and standard deviations estimated from the generalized mixed effects model. The star (*) indicates a significant interlimb difference (p<0.05).

**Figure 5:**
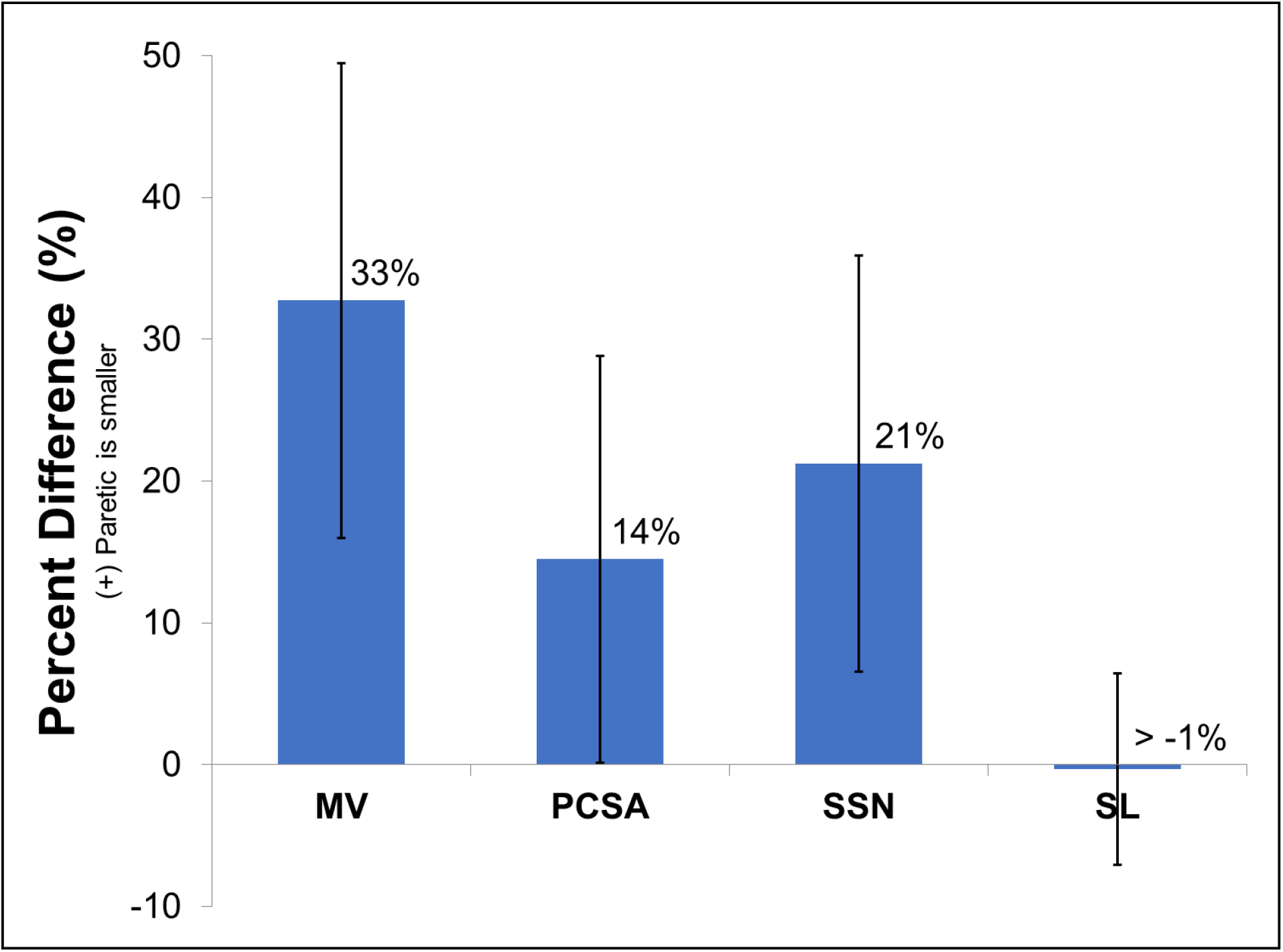
Interlimb Muscle Differences Post-Stroke. Bar graph showing the percent difference in muscle architectural (sarcomere length “SL” and muscle volume “MV”) and functional parameters (serial sarcomere number “SSN” and physiological cross sectional area “PCSA”) estimated from the general linear mixed-effects model for all stroke participants who had smaller muscle volume on the paretic biceps. A positive percent difference indicates that the paretic parameter is smaller than the non-paretic side. Error bars represent one standard deviation from the mean of the percent difference across the subjects.

## DISCUSSION

This study aimed to quantify *in vivo* differences in muscle architecture parameters between the paretic and non-paretic biceps brachii of individuals with chronic hemiparetic stroke. Most notably, this comprehensive, multi-scale study found fewer sarcomeres in series in the paretic muscle compared to the contralateral side. In 7 of 8 stroke participants, we observed strikingly smaller muscle volumes on the paretic side. However, the corresponding deficit in PCSA of the paretic biceps, the architectural parameter that predicts the maximum isometric force the muscle can generate with full activation(25), was more modest. In these seven stroke participants, the fact that each had fewer serial sarcomeres partially explains the smaller paretic muscle volume. We quantified the same architectural parameters in age-range matched individuals who had not undergone a stroke; we found no substantial or significant interlimb differences in any muscle architecture parameter, suggesting the interlimb differences we observed were adaptations associated with chronic stroke rather than natural interlimb variability. Thus, the architectural parameters suggest a functional re-organization of the muscle. Specifically, shorter optimal fascicle lengths are generally understood to indicate a proportional decrease in the width of the muscle’s isometric force-length curve^21,24,43^, or – more explicitly – a proportional decrease in the absolute range of lengths over which the muscle can generate active force. Given this adaptation in length, the loss of muscle volume we observed in 7 of the 8 paretic limbs was not a direct measure of the loss in the muscle’s force-generating capacity (Fig. 5). The substantial decrease in serial sarcomere number that occurs when a muscle is held at a joint posture which places it at a shortened length was first reported in classic limb immobilization studies in animal models(18, 27, 29); it is widely assumed to be a fundamental muscle adaptation process. We now provide the first direct evidence of this phenomenon in living human subjects.

The most complete demonstration of in vivo muscle adaptation that accompanies chronic length changes has been via animal models involving limb immobilization(18, 27–29). The main difference between the adaptations observed in our study compared to these studies of limb immobilization is the magnitude of the observed adaptation. The decrease in SSN we observed in the paretic biceps post stroke was less substantial than the loss of serial sarcomeres observed following immobilization at a shortened muscle-tendon length in animal models (~22% vs ~40%). The prime differences between individuals post-stroke and animal immobilization studies, which may explain this difference in magnitude, is that post-stroke individuals tend to disuse their paretic limb, but still have active and passive joint motion whereas animal immobilization studies eliminated movement entirely. In our stroke participants, the studied muscle also receives altered neural inputs; the animal studies did not involve a neural injury.

In human subjects, only a single previous study has ever measured both sarcomere lengths and fascicle lengths in the same muscle under conditions that are generally assumed to result in a loss of serial sarcomeres (i.e., in a muscle that has been chronically placed in a shortened position)^32^. This previous work evaluated the soleus muscle in children with cerebral palsy who were undergoing surgery to address equinus contractures. However, because of the invasive methods required to measure sarcomere lengths before the development of second harmonic generation microendoscopy, measures of sarcomere length were obtained intraoperatively. Reasonably, these data could not also be collected from typically developing children. Thus, despite the valuable data obtained from the clinical population, serial sarcomere number in the chronically shortened soleus muscle could not be compared to direct measures of the same parameters obtained in a control population. Despite this limitation, the intraoperative data provided evidence that the sarcomere lengths in the chronically shortened soleus were extremely long compared to optimal sarcomere length, even in an extremely plantarflexed limb posture. This finding was surprising because the results from the immobilization studies in animal models suggest that muscle “re-optimizes” such that optimal length occurs in the limb position of immobilization. Importantly, this result in the soleus replicated a previous finding in children with CP undergoing surgery for wrist contracture. In this case, sarcomere lengths for the wrist flexor in a flexed wrist position were shown to be much longer compared to typically developing children undergoing surgery to treat radial nerve palsy (3.48μm vs 2.41μm)(50). Unlike the results in children with CP, we did not observe systematically longer sarcomeres in the paretic muscles of individuals with chronic hemiparetic stroke (Fig. 2B). We expect that a critically important difference between muscle adaptation following stroke and CP is that stroke occurs in a fully developed system. Similar to the biceps brachii in our population, the affected muscles in the CP studies were chronically in a shortened positioned due to the primary neural impairments. However, it has been posited that while chronically shortened CP muscle loses sarcomeres in series as would be expected from the classic immobilization studies, a malfunctioning sensing system within the muscles prevents the addition of serial sarcomeres during bone growth, resulting in the abnormally long sarcomeres(30). Thus, the additional physiological process of bone growth in children with CP confounds direct comparison with muscle adaptation that occurs in adults with stroke.

In general, our work provides the most direct confirmation in humans to date that chronic impairments that lead to disuse and place a muscle in a shortened position are associated with the loss of serial sarcomeres. In the context of the literature discussed above, we note that this adaptation process also seems to be moderated by the presence of confounding factors (i.e. bone growth (CP), altered neural input, disuse resulting in a reduced use of available range of motion, etc.). Such confounds are common *in vivo* following disease, neural injury, and clinical interventions such as surgery or immobilization. With the comprehensive multiscale imaging techniques utilized in this study, adaptation of muscle structure in the context of confounding factors can now be better explored and more fully understood in humans, which is necessary for the development of more targeted interventions that seek to improve outcomes.

The substantial decrease in serial sarcomere number found in the paretic biceps brachii likely amplifies motor impairments which stem from stroke-induced neural impairment. Decreased voluntary neural drive (weakness or paresis)(7, 8), increased involuntary neural drive (hypertonicity), and abnormal muscle coactivation patterns(9) occur following damage of the corticofugal motor pathways due to a stroke. In addition, it is clear that the stroke induced neural deficits result in altered use of the contralesional or paretic limbs. Particularly notable is the difficulty (and in severe cases impossibility) of coordinated extension of the upper limb to reach and grab an object which is some distance from the body(12, 51). In this study we find that the muscle structure itself has been altered via a loss of serial sarcomeres in the paretic biceps brachii muscle. Functionally, for the individuals in our study who had survived a stroke, this means that the paretic muscle has a narrower range of lengths and joint angles over which it can produce active force. Reduction in serial sarcomeres resulted in an increase in the muscle’s passive resistance to stretch in animal models of limb immobilization(29). For the biceps, this would exacerbate neurally-driven motor impairments that diminish the ability to extend the elbow, in the presence of significant elbow extensor weakness, to reach for an object away from their body.

Results from our study are reasonable in the context of previously reported in vivo studies which independently measure either fascicle length, muscle volume, or sarcomere length. Specifically, muscle fascicle lengths in the paretic biceps brachii in this study were on average 20.6% shorter than the non-paretic side. Previous studies performed in elbow flexors report similar, substantial decreases in fascicle length in extended joint postures (18.6% decrease in biceps brachii at 25° elbow flexion (46),15% decrease in brachialis at 10° elbow flexion(45)). Studies of muscle volume differences between paretic and non-paretic limbs are variable. On average, our muscle volume differences (29%) are in the same direction (paretic muscle is smaller) and of slightly greater magnitude than other upper limb studies (no difference to 25% difference(40, 47)) and lie within the range of lower limb difference (no difference to 33%^35,49^). The exclusion of intramuscular fat in our measure of muscle volume is relatively novel and may explain our slightly higher percent differences. Measurements of biceps brachii sarcomere lengths have only been obtained in vivo in a single study that enrolled 4 individuals with chronic hemi-paretic stroke. Although there was not a significant difference in the mean sarcomere length between limbs among the stroke participants we studied, similar to this previous work, we found that some individuals had shorter sarcomere lengths in their paretic limb while others had longer(49). Notably, independently measured muscle anatomical parameters (particularly sarcomere lengths and non-normalized fascicle lengths) are difficult to directly compare between studies as they are sensitive to the limb posture chosen for testing(52); muscle fiber and sarcomere lengths change with joint position.

There are various limitations to this study. While it provides novel *in vivo* evidence of comprehensive muscle architecture changes following chronic hemiparetic stroke, more participants, a larger number of muscles studied in more joint positions, and inclusion of factors that quantify the extent participants use their upper limbs (i.e. passive and active range of motion, elbow joint resting posture, amount of voluntary use of limb, etc.) would broaden our knowledge of muscle adaptation post brain injury. We would expect that altered tendon properties or increased tendon length(53) could accompany the paretic muscle adaptation we observed, but we did not include these measures in this study. This study is limited in that we did not directly measure stiffness or changes in muscle or tendon properties which may further elucidate muscle-tendon adaptation post-stroke.

Beyond the addition of the first comprehensive measurements of in vivo muscle architecture for the investigation of muscle plasticity to stroke-induced neural deficits, this study demonstrates the need for such comprehensive in vivo studies and provides insight for the design of therapeutic interventions for stroke survivors. Without the combination of fascicle and sarcomere lengths measured at the same joint angle with quantification of muscle volume, ambiguity would remain in the functional impact of these individual muscle parameters. Many prior studies in stroke and other populations demonstrate interlimb differences in fascicle length without normalizing to sarcomere length. With the addition of sarcomere length, we were able to explicitly demonstrate that the paretic biceps brachii muscle has fewer serial sarcomeres. Our finding leads us to conclude that stroke-induced neural deficits, which lead to altered input and disuse of the contralesional limb, ultimately change the basic biceps brachii muscle architectural parameters in a way which may amplify functional impairments.

## METHODS

### Participants

Measurements of sarcomere length, fascicle length, and muscle volume of the biceps brachii were obtained using in vivo imaging techniques in both arms of twelve participants; eight participants with chronic hemiparetic stroke (3 female/5 male, 60 ± 9 yrs, Fugl-Meyer 26 ± 9, 13 ± 10 yrs post-stroke) and four participants with no history of musculoskeletal or neurological diseases or injuries to the upper limb (2 female/2 male, 62 ± 6 yrs). Fugl-Meyer Assessment scores reported in this study were performed by a licensed physical therapist prior to experimentation. All the individuals who participated in this study provided informed consent prior to experimentation; Northwestern University’s Institutional Review Board approved this study’s procedures.

### Sarcomere length

Sarcomere lengths of both arms were acquired with the participants seated in a comfortable chair. The arm being imaged was secured at 85° shoulder adduction, 10° horizontal shoulder flexion, 25° of elbow flexion, mid pro-supination, and 0° wrist and finger flexion (Fig. 6). Joint posture was verified by goniometric measurement. A microendoscopic system (Zebrascope, Enspectra Health (previously Zebra Medical Technologies), Mountain View, CA) which consists of a laser (class IV, output power > 500mW, center wavelength 1030 nm), microscope, and microendoscopic probe, was used to image sarcomeres in vivo. This system utilizes the second-harmonic generation optical technique to capture the intrinsic striation pattern of sarcomeres(36, 49).

**Figure 6:**
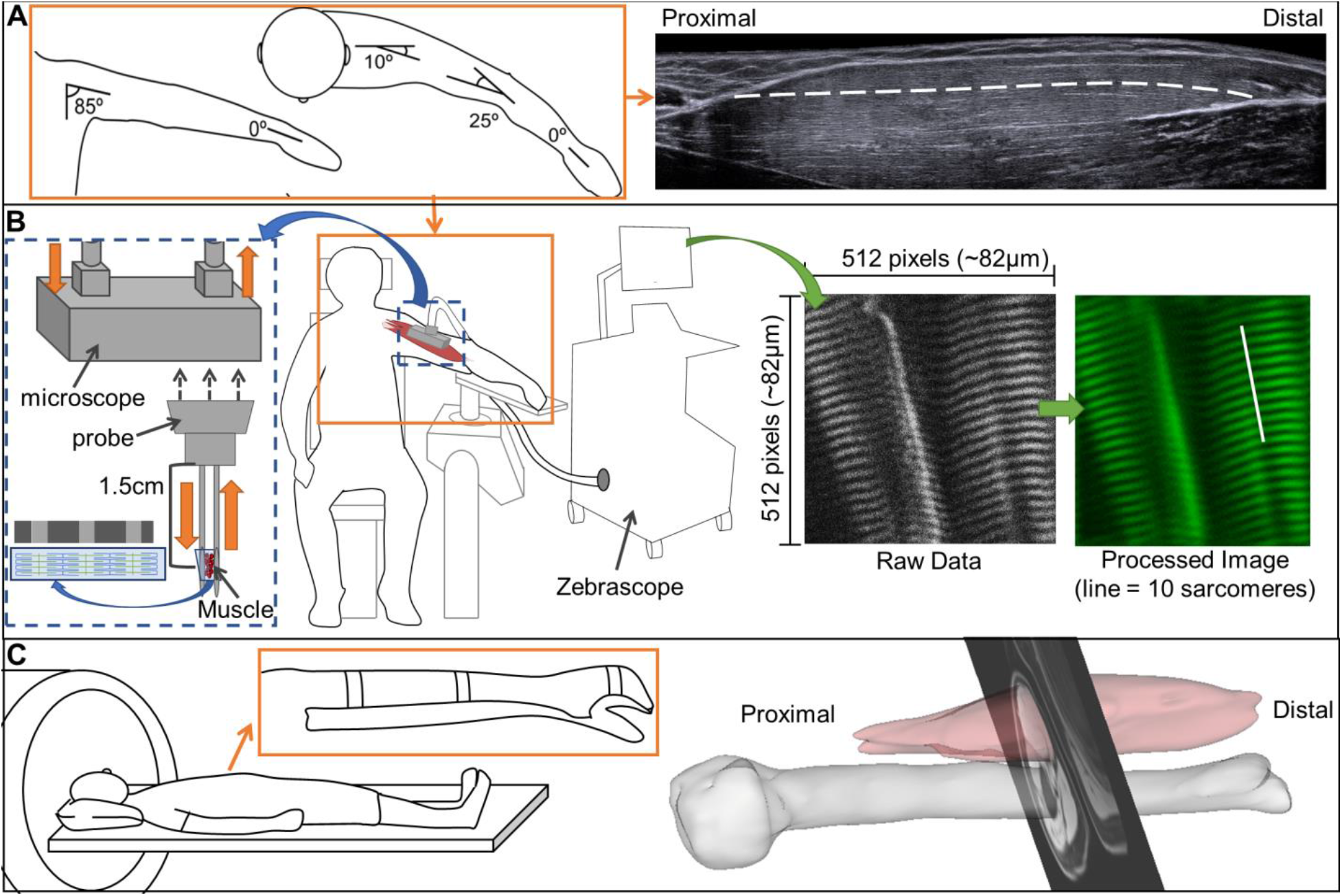
Illustration of experimental set up for muscle architecture measures. **A.** Arm posture used for measurement of fascicle length using ultrasound (right) and for sarcomere length (see B). The white dashed line overlaid on the ultrasound image demonstrates a muscle fascicle. **B.** The illustration shows the Zebrascope which utilizes second harmonic generation microendoscopy to capture the natural striation pattern of sarcomeres in vivo. The Biodex chair and arm fixture (middle) used for measurement of sarcomere and fascicle lengths (see A). The microscope is blown up on the left (Blue dotted box) to show the flow of laser light through the microscope (orange arrows). Below the microscope is a graphic of the probe which is inserted into the bicep brachii muscle. Between the two needles of the probe at 1.5cm the laser light interacts with myofibrils and the striation pattern of the sarcomeres can be captured. To the right, a raw image which would be seen during image collection is shown. After post-processing, sarcomere length is measured from the processed image which is showing the length of 10 sarcomeres in series (white line). **C.** Participant in the supine positioning (left) in the MRI bore. Orange inset shows the splinting of the arm to reduce artifacts due to hand and arm movement. (Right) 3D rendering of the biceps brachii muscle and humerus bone with a single MRI slice superimposed.

A sterile microendoscopic probe was inserted into the long head of the biceps brachii. The probe consisted of two 1.8cm long, 20-gauge needles with beveled tips (Fig. 6B); one needle with a transmitting lens used to excite the muscle tissue and one needle housing a receiving lens to capture the reflection of the signal after it has interacted with tissue. Ultrasound and palpation techniques were used, prior to insertion, to verify placement of the probe at mid-belly of the muscle with the probe’s optical lenses aligned parallel to the fascicle direction. A spring-loaded injector was used to rapidly insert the probe, minimizing pain and improving precision of probe placement. The microscope was attached to the microendoscopic needle for imaging. Images with a field-of-view of 82μm by 82μm were collected at 1.9Hz for approximately 2-5 minutes. The image produced from the microendoscopic system captures the A-bands (myosin protein) of the sarcomeres and enables direct measurement of sarcomeres from the resulting striation pattern(49). Surface EMG (Bagnoli-16 system, Delsys Inc., Boston, MA) of the biceps brachii was obtained simultaneously using a custom written MATLAB script. Baseline EMG activity was collected for 10 seconds with the needle inserted and participant relaxed. Muscle activation was visually monitored during data collection and analysis was performed offline.

### Fascicle length

Fascicle length measurements of the long head of the biceps brachii in both arms of all participants were obtained using extended field-of-view ultrasound (EFOV-US) under the same conditions (same joint posture, passive muscle) as sarcomere length measures. The extended field-of-view technique involves sweeping the ultrasound probe along the length of the muscle as sequential B-mode ultrasound images are acquired and stitched together to form a single composite image with a field-of-view longer than the ultrasound probe’s aperture (±60 cm)(54). This method has been demonstrated to be accurate and reliable for measurement of fascicle length in different individuals and muscles(52, 55). Approximately 10 qualitatively good images were captured per arm. EFOV-US images (Acuson S2000, linear array transducer 18L6, SieScape, Siemens Medical Solutions USA, Inc., Mountain View, CA) and surface EMG (Bagnoli-16 system, Delsys Inc., Boston, MA) of the biceps brachii were simultaneously recorded (Spike, Power1401, and Micro1401-3, Cambridge Electronic Design Limited, Cambridge, England).

### Muscle Volume

To determine the volume of the biceps brachii muscle, excluding intramuscular fat, the Dixon method, a fat suppression MRI sequence, was implemented on both upper limbs of all participants (3D GRE, TR = 7ms, flip angle = 12°, matrix size = 256 x 304, slice thickness = 3mm, TE of 2.45ms and 3.68ms)(47). As increases in intramuscular fat within muscle has been demonstrated in the lower limb of stroke participants(41) and patients with other pathologies (i.e whiplash(56), spinal cord injury(57)) correcting muscle volumes for the amount of intramuscular fat is necessary to avoid potential overestimation of volume of muscle. The participants were lying supine in a 3T MRI (Area, Siemens Medical Solutions USA, Inc., Mountain View, CA) scanner with their arm as close to the center of the scanner as possible. To minimize participant movement during scanning, the lower arm was splinted using an orthosis.

### Data analysis

#### Sarcomere length

Image sequences obtained from the microendoscopic system were post-processed offline using a script provided with the ZebraScope by Enspectra Health (previously Zebra Medical Technologies, Mountain View, CA). With this script, the raw image sequence was first combined into a multipage Tiff. Then, a Fast Fourier Transform (FFT) cleared the edges and vertical center which contained only noise; the transformed data were symmetrically, low pass, Gaussian filtered. Within the script, we set the frequency bounds to filter out all frequencies which would yield unphysiologic sarcomere length values, specifically, values smaller than 2μm or larger than 5μm. This removed all images without sarcomeres (without frequencies in the 2-5 μm range) from the multipage Tiff sequence. White noise in the FFT was calculated and removed. To determine mean sarcomere length from each processed image, the peak frequency of a least squares fit of a Gaussian was calculated. The final outputs of the image processing script for each arm and each participant were mean sarcomere length and standard deviation, calculated using all processed images that were not excluded by the specified frequency bounds. At this point, we used our own custom-written MATLAB code to further exclude images that were collected when the biceps EMG signal was 3 standard deviations above the resting baseline EMG. Thus, the mean sarcomere lengths and standard deviations that we report for each biceps brachii were calculated from the processed image data, further restricted by the synced EMG data to only include images that were collected under passive conditions.

#### Fascicle Length

EFOV-US images were exported as DICOM images and measurements of fascicle length were made offline using the segmented line tool in ImageJ (ImageJ with Fiji, version 1.51h, Wayne Rasband, National Institutes of Health, Bethesda, MD(58)). Measurements were made on 3 images per arm per participant. The experimenter selected the 3 images which best captured the entire muscle and had visible fascicles which extended from central tendon to aponeurosis. Four fascicles were measured per image. Mean fascicle length was calculated across the 3 images (3 images X 4 fascicles per image = 12 fascicles)(46, 52).

#### Muscle Volume

Manual segmentation of the biceps brachii (long and short head) muscle was performed using Analyze12.0 (AnalyzeDirect, Overland Park, KS). To calculate the volume of muscle without intramuscular fat (*V_m-f_*), the following equations were implemented:

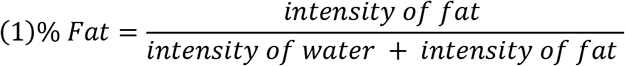

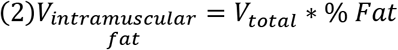

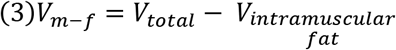

Where *V_total_* is the total volume segmented from the MRI images. Measurements of biceps muscle volume without fat were made by two different raters (rater 1 n = 7, rater 2 n=5). Segmentation and calculation of muscle volume without intramuscular fat (*V_m-f_*) has been shown to be reliable within and across raters(47).

#### Calculation of Functional Parameters

With the quantification of sarcomere length, fascicle length, and muscle volume, optimal fascicle length (*OFL*), serial sarcomere number (SSN), and physiological cross-sectional area (PCSA), were calculated for both arms of all participants using the following equations.

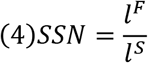

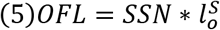

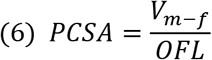

Where 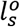 is 2.7 *μm* or optimal sarcomere length(33), and *l^F^* (fascicle length), *I^s^* (sarcomere length), and *V_m-f_* (volume of muscle without fat infiltration) were measured in this study. For statistical analysis involving equations (4–6), average fascicle length and all measures of sarcomere length were utilized.

### Statistical Analysis

To determine if there were significant interlimb differences in any of the muscle architecture parameters or calculations, generalized linear mixed-effects models were implemented (SAS 9.4, SAS Institute Inc., Cary, NC). Each model had one of the parameters or calculations as the outcome variable. Whether or not the participant had a stroke was a fixed effect in the model. Within-subject correlation and the correlation between paretic and non-paretic (or dominant non-dominant) arms were modeled as random effects. A significant difference between limbs was present if the p-value was less than 0.05 for all models. To determine if there is a linear relationship between the percent difference in SSN or OFL in the participants studied with stroke and their clinical function score (Fugl-Meyer assessment), a linear regression was performed.

An a priori power analysis was conducted to determine sufficient sample size to test the hypothesis that participants with stroke have interlimb differences in optimal fascicle length which are not present in individuals who have not had a stroke. The analysis indicated that with 8 participants with stroke and 4 participants without stroke and an effect size greater of 1.85, a power greater than 0.8 would be achieved. This effect size was established from interlimb differences in fascicle and sarcomere length obtained from two previous studies performed on different sets of individuals(46, 49). The effect size from our data was verified to exceed the effect size from the a priori analysis.

## AKNOWLDGEMENTS

We would like to thank the study participants, Vikram Darbhe for assistance in data collection, and Dr. Masha Kocherginsky and Liqi Chen for aid with statistical analysis. We would also like to thank Sabeen Adamani for assistance in equipment set up and troubleshooting and Zebra Medical Technologies (now Enspectra Health) for their support with data collection and image processing. This work is supported by the National Science Foundation (NSF) Graduate Research Fellowship Program under Grant No. DGE-1324585, as well as National Institutes of Health (NIH) R01 D084009. Any opinions, findings, and conclusions or recommendations expressed in this material are those of the authors and do not necessarily reflect the views of the NSF or NIH.

## Notes

### Competing Interest Statement

The authors have declared no competing interest.

